# A Phytochrome B-PIF4-MYC2 Module Tunes Secondary Cell Wall Thickening in Response to Shaded Lighting

**DOI:** 10.1101/2021.10.11.463962

**Authors:** Fang Luo, Qian Zhang, Hu Xin, Hongtao Liu, Hong-Quan Yang, Monika S Doblin, Antony Bacic, Laigeng Li

## Abstract

Secondary cell walls (SCW) in stem xylem cells provide mechanical strength and structural support for growth. SCW thickening is light- regulated and varies under different light growth conditions. Our previous study revealed that blue light enhances SCW thickening through the activity of MYC2 directed by CRYPTOCHROME1 (CRY1) signaling in stem xylary fiber cells. In this study, we demonstrate that the low ratio of red: far-red light (R:FR) of the shaded light condition inhibits SCW thickening in the inflorescence stem of *Arabidopsis*. Phytochrome B (PHYB) plays a dominant role in perceiving the R:FR balance. Under white and red-light conditions, *phyB* mutants display thinner SCWs in xylary fibers, but thicker SCWs are deposited in the PHYTOCHROME INTERACTING FACTORS (PIFs) quadruple mutant *pif1pif3pif4pif5* (*pifq*), suggesting involvement of the PHYB-PIFs signaling module in regulating SCW thickening. Interaction of PIF4 with MYC2 affects MYC2 localization in nuclei and inhibits its transactivation activity on the *NST1* promoter. Shade conditions mediate the PIF4 interaction with MYC2 to regulate SCW thickening. Genetic analysis confirms that the regulation of SCW thickening by *PIFs* is dependent on *MYC2* function. Together, these data reveal a molecular mechanism for the effect of shaded light inhibition on SCW thickening in stems of *Arabidopsis*.

## Introduction

In higher plants, all cells are encased in a primary cell wall laid down during cell elongation that is flexible (to allow growth) yet possesses sufficient tensile strength to withstand the turgor pressure that drives growth. The primary cell wall defines the shape of a plant cell and is important for communication between plants and their environment(Doblin et al., 2010). Some types of specialized cells such as fiber and vessel cells in stem xylem deposit a rigid secondary cell wall (SCW) inside the primary cell wall after cell elongation has ceased, which provides plants with the mechanical strength to withstand enormous compressive forces and the capacity to transport water to aerial organs(Zhong and Ye, 2015). During plant growth, the plant body is built and structured primarily by conversion of photosynthetic products into SCW. The main components of lignified SCW include cellulose, hemicelluloses and lignin, with deposition of lignin being a sign of SCW thickening. Expression of SCW biosynthesis genes is controlled by a hierarchy of transcriptional regulatory networks. *SND1*/*NST1* and *VND6*/*VND7* are key regulators at the top tier of the regulatory network that specifically control SCW formation in fiber and vessel cells, respectively, in *Arabidopsis*(Zhong et al., 2010; Taylor-Teeples et al., 2015).

In addition to developmental signals, various external environmental cues, including light, water and temperature, affect SCW formation(Le Gall et al., 2015). Light induces a range of effects on plant cell wall formation(Le Gall et al., 2015). For example, when grown under blue light, *Arabidopsis* deposits a mechanically strengthened inflorescence stem due to a thickening of the SCW of fiber cells. It was demonstrated that the blue light signal induces *MYC2* expression which activates *NST1* expression by binding to its promoter, leading to an enhancement of SCW thickening(Zhang et al., 2018a). However, under shaded light conditions with a lower ratio of red to far-red (R:FR) light, plants exhibit an increase of cell elongation and a subsequent reduction in their SCW thickening(Sasidharan et al., 2008; Sasidharan et al., 2010; Casal, 2012; Pedmale et al., 2016; Wu et al., 2017). The molecular mechanism underlying this reduction of SCW thickening caused by shaded light conditions remains unknown.

The red/far-red photoreceptor, phytochrome B (PHYB) exists in two forms that are reversibly interconvertible by perceiving red and far-red light(Quail, 1991). The Pr form of PHYB absorbs red light to rapidly revert to the Pfr form which absorbs far-red light to return to the Pr form(Franklin, 2008). In response to the R:FR ratio signal, interconversion of the PHYB Pr-Pfr forms activates downstream molecular pathways to regulate induction of seed germination, seedling de-etiolation, shade avoidance, and floral initiation(Franklin and Quail, 2010; Strasser et al., 2010).

Red light activates PHYB to interact with PHYTOCHROME INTERACTING FACTORs (PIFs, mainly a quartet of members: PIF1, PIF3, PIF4, and PIF5), leading to their degradation(Bauer et al., 2004; Monte et al., 2004; Al-Sady et al., 2006; Shen et al., 2007; Lorrain et al., 2008), whereas far-red light inactivates PHYB and stabilizes PIFs, inducing stem elongation and other morphogenesis processes(Hornitschek et al., 2012; Leivar et al., 2012). However, how PHYB responds to the R:FR ratio in regulation of the SCW thickening is largely unknown.

The transcription factor (TF) MYC2 is considered a transcriptional regulatory “hub” interconnecting a variety of biological processes(Kazan and Manners, 2013). MYC2 interacts with different TFs to integrate the crosstalk among different signaling pathways including jasmonate (JA)-mediated pathogen defences, abscisic acid (ABA), ethylene, gibberellic acid (GA) and light signals(Chen et al., 2012; Hong et al., 2012; Song et al., 2014). The blue light signal upregulates expression of *MYC2* which then activates the *NST1*–mediated SCW thickening by direct binding to the *NST1* promoter(Zhang et al., 2018a). Here, we show that the low R:FR ratio of a shaded light condition inhibits SCW thickening in *Arabidopsis* inflorescence stems. Our analyses indicated that the R:FR light is perceived by PHYB to alter the status of PIF4, which act as a direct regulator of MYC2 to modulate SCW thickening. This study reveals a molecular pathway of the SCW thickening in response to R:FR ratio light conditions.

## Results

### Light R:FR ratio affects SCW thickening in the inflorescence stem of *Arabidopsis*

As our previous study showed that a blue light signal enhances SCW thickening in *Arabidopsis* inflorescence stems(Zhang et al., 2018a), we were interested in further examining the effect of light with a lower ratio of R:FR, which imitates shaded light conditions, on SCW thickening during inflorescence stem growth. First, wild-type (WT) plants were grown in normal white light (WL) conditions to bolting. When the inflorescence stem started to grow, plants were transferred to light conditions with a different R:FR ratio and the growth of the inflorescence stem monitored. The inflorescence stem grew much faster (**Figure 1, A and B)** and had a significantly lower tensile strength (**Figure 1C**) under white light supplemented with far-red light (FR) (WL+FR) compared to white light (WL). This shows that the low R:FR ratio condition affected both inflorescence stem growth and mechanical strength. Anatomical analyses of the stem structure revealed that the SCW thickness in fiber cells was significantly decreased, but vessel cell SCW thickness showed little difference (**Figure 1, D and E**). Expression of the SCW thickening marker genes (*NST1*, *SND1*, *4CL1* and *IRX8*)(Lee et al., 1997; Zhong et al., 2006; Mitsuda et al., 2007; Hao et al., 2014) were down-regulated under low R:FR conditions (**Supplemental Figure S1**). Content of lignin and crystalline cellulose, the typical SCW components, was significantly lower under low R:FR condition (**Figure 1, F and G**). These data suggest that the additional far-red light promotes stem elongation and inhibits SCW thickening in stem fiber cells.

**Figure 1.**
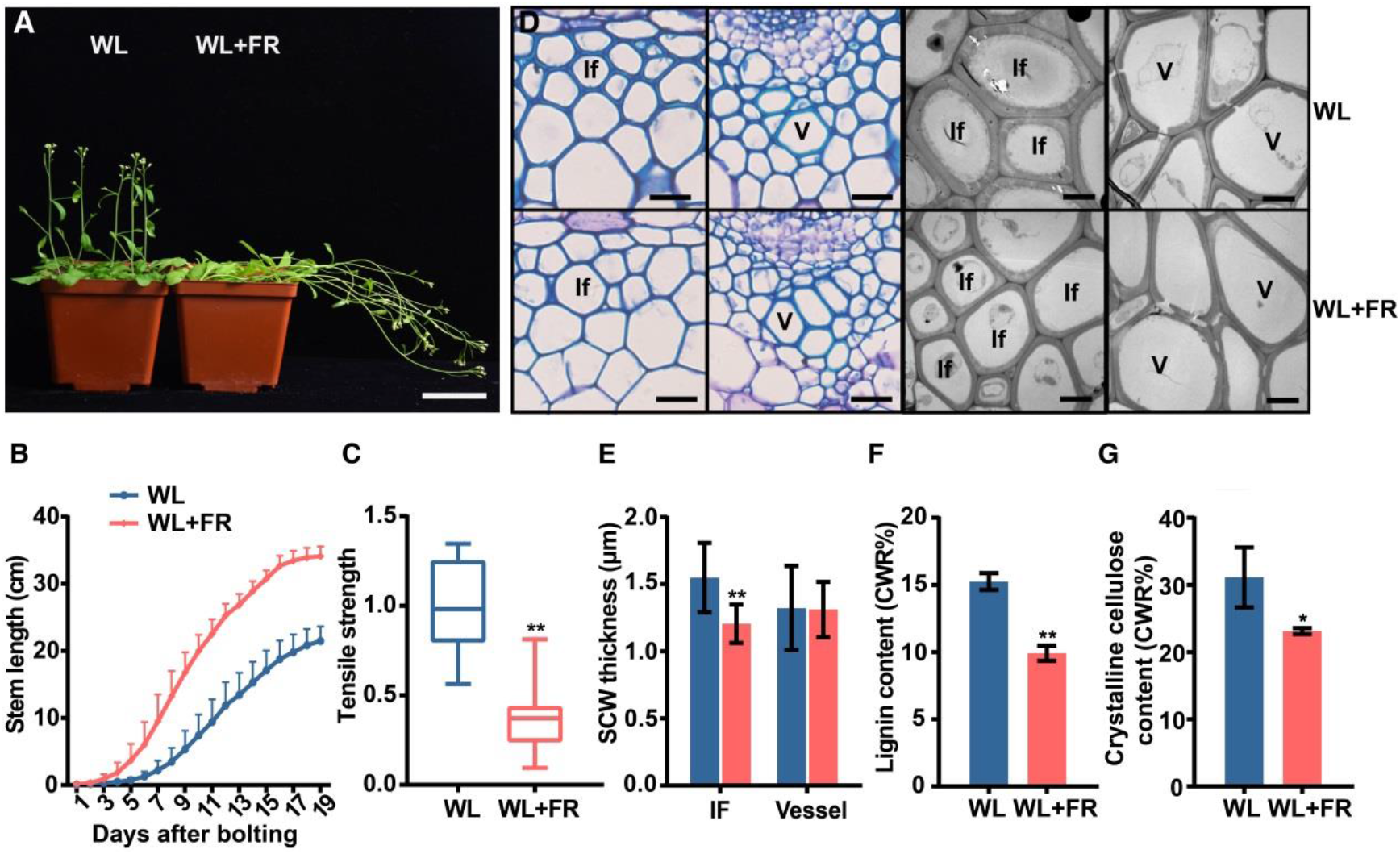
Shaded light inhibits SCW thickening in inflorescence stem. **A,** Growth of *Arabidopsis* inflorescence stems in WL and WL+FR conditions. WL: white light. FR: far red. Scale bar = 5 cm. **B,** Elongation of inflorescence stems in WL and WL+FR conditions during growth. N = 9, mean ± SD. **C,** Tensile strength of the inflorescence stems. Student’s t test (**P < 0.01) was used for statistical analysis, n = 18, mean ± SD. **D,** Cross sections of the inflorescence stem grown under different light conditions (WL and WL+FR) visualized under the light microscope (after Toluidine blue staining; LHS panels) and the transmission electron microscope (RHS panels). If: Interfascicular fiber cell. V: Vessel cell. LHS panels: scale bar = 20 μm, RHS panels: scale bar = 5 μm. **E,** Measurements of SCW thickness in the interfascicular fiber cells in (**D**). From three biological replicates, more than 10 cells were measured per biological replicate. Student’s t test (**P < 0.01) was used for statistical analysis, mean ± SD. **F,** Lignin content in inflorescence stems of the plants grown in different R:FR conditions. Student’s t test (*P < 0.05) was used for statistical analyses, n = 3, mean ± SD. **G,** Crystalline cellulose content in inflorescence stems of the plants grown in different R:FR conditions. Student’s t test (*P < 0.05) was used for statistical analyses, n = 3, mean ± SD.

### PHYB and PIFs are involved in regulating cell elongation and SCW thickening in *Arabidopsis* inflorescence stem

R/FR light is perceived by the photoreceptor PHYB which induces a series of responses through PIF proteins(Reed et al., 1993; Pham et al., 2018). To dissect the genetic base of the light R:FR ratio effect on SCW thickening, we analyzed the inflorescence stem growth of *phyB* and quadruple *pif1pif3pif4pif5* (*pifq*) mutants. *phyB* mutants displayed a lodging phenotype and grew longer inflorescence stems than wild-type (WT), while *pifq* mutants showed erect growth with shorter inflorescence stems compared to WT (**Figure 2, A and B; Supplemental Figure S2A**). Stem elongation growth was determined by measuring the distance between two markers on the inflorescence stem during its growth process. The elongation growth was increased in *phyB* but decreased in *pifq* mutant plants (**Figure 2C**). The inflorescence stem mechanical properties, measured as tensile strength, were also significantly impacted: *phyB* inflorescence stems had decreased tensile strength, whereas the tensile strength of *pifq* mutant inflorescence stems was increased (**Figure 2G**). To analyze the cell length and cell wall structure in the inflorescence stem, the stem was disintegrated to measure the length of xylem fibers which are the predominant cell type in mature inflorescence stems. The fiber cells of *phyB* plants were longer whereas those of *pifq* plants were shorter than those in WT (**Figure 2D; Supplemental Figure S2B**).

**Figure 2.**
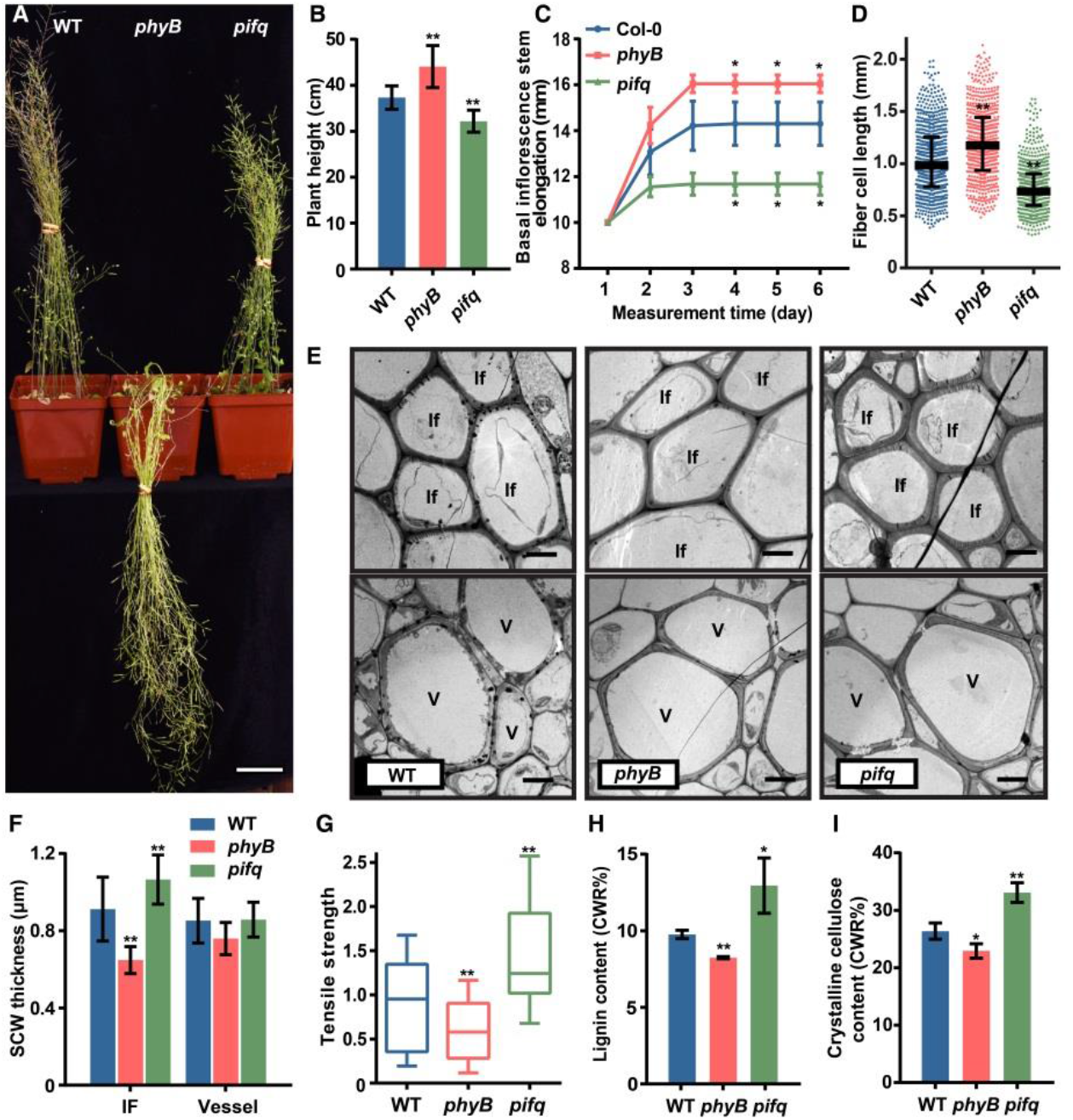
PHYB and PIFs regulate SCW thickening in fiber cells. **A,** Plants were grown in white light (WL) at 8-weeks old. Scale bar = 5 cm. **B,** Measurements of plant height in (**A**). Student’s t test (**P < 0.01) was used for statistical analyses, n = 20, mean ± SD. **C,** Inflorescence stem elongation. The stem was marked with two points at the basal region and the distance between the two points was measured every day during growth. Student’s t test (*P < 0.05) was used for statistical analyses, n = 3, mean ± SD. **D,** Fiber cell length measured in disaggregated fiber cells. Data were from three biological replicates and more than 200 cells were measured per biological repeats. Student’s t test (**P < 0.01) was used for statistical analysis, mean ± SD. **E,** Transmission electron micrographs of inflorescence stem cross-sections. If: Interfascicular fiber cell. V: Vessel cell. Scale bar = 5 μm. **F,** Measurements of SCW thickness in the interfascicular fiber cells and vessel cells in (**E**). Data were from three biological replicates and more than 5 cells were measured per biological replicate. Student’s t test (**P < 0.01) was used for statistical analysis, mean ± SD. **G,** Tensile strength measurements of the inflorescence stem. Student’s t test (**P < 0.01) was used for statistical analysis, n = 16, mean ± SD. **H,** Lignin content in inflorescence stems. Student’s t test (**P < 0.01, *P < 0.05) was used for statistical analyses, n = 3, mean ± SD. **I,** Crystalline cellulose content in inflorescence stems. Student’s t test (**P < 0.01, *P < 0.05) was used for statistical analyses, n = 3, mean ± SD.

Meanwhile, cross-sections of the stem showed a difference in the cell wall thickness in interfascicular fiber cells (**Figure 2F; Supplemental Figure S2C**). Compared to WT, cell wall thickness in interfascicular fiber cells was decreased in *phyB* but increased in *pifq* plants, whereas the thickness of vessel cells was unaltered (**Figure 2, E and F**). Furthermore, both the lignin and crystalline cellulose content was decreased in *phyB* but increased in *pifq* plants (**Figure 2, H and I,** respectively).

In parallel with the knock-out mutant analyses, *Arabidopsis* plant overexpressing *PHYB* (*PHYB-OE*) and *PIF4* (*PIF4-OE*) were also generated, and their inflorescence stem properties analyzed. *PHYB-OE* transgenics had shorter and stronger (increased tensile strength) inflorescence stems while those of *PIF4-OE* plants were thinner and weaker (**Supplemental Figure S3**; **Figure 3C**). Examination of stem cross-sections indicated that *PHYB* overexpression resulted in thicker cell walls in xylary interfascicular fiber cells while *PIF4* overexpression led to thinner fiber cell walls relative to WT (**Figure 3, A and B)**. Lignin and crystalline cellulose content was increased in the *PHYB-OE* inflorescence stems but decreased in *PIF4-OE* stems (**Figure 3, D and E)**. In contrast, the cell wall thickness of vessel cells was largely unaffected in *PHYB-OE* but was decreased in *PIF4-OE* plants (**Figure 3, A and B**). Transcriptional analyses of the SCW thickening genes (*NST1*, *SND1*, *4CL1* and *IRX8*) demonstrated that the expression of these genes was upregulated in *PHYB-OE* plants but suppressed in *PIF4-OE* plants (**Figure 3F**). These results indicate that SCW thickening is positively regulated by *PHYB* but negatively regulated by *PIF*s in inflorescence stem xylary fibers.

**Figure 3.**
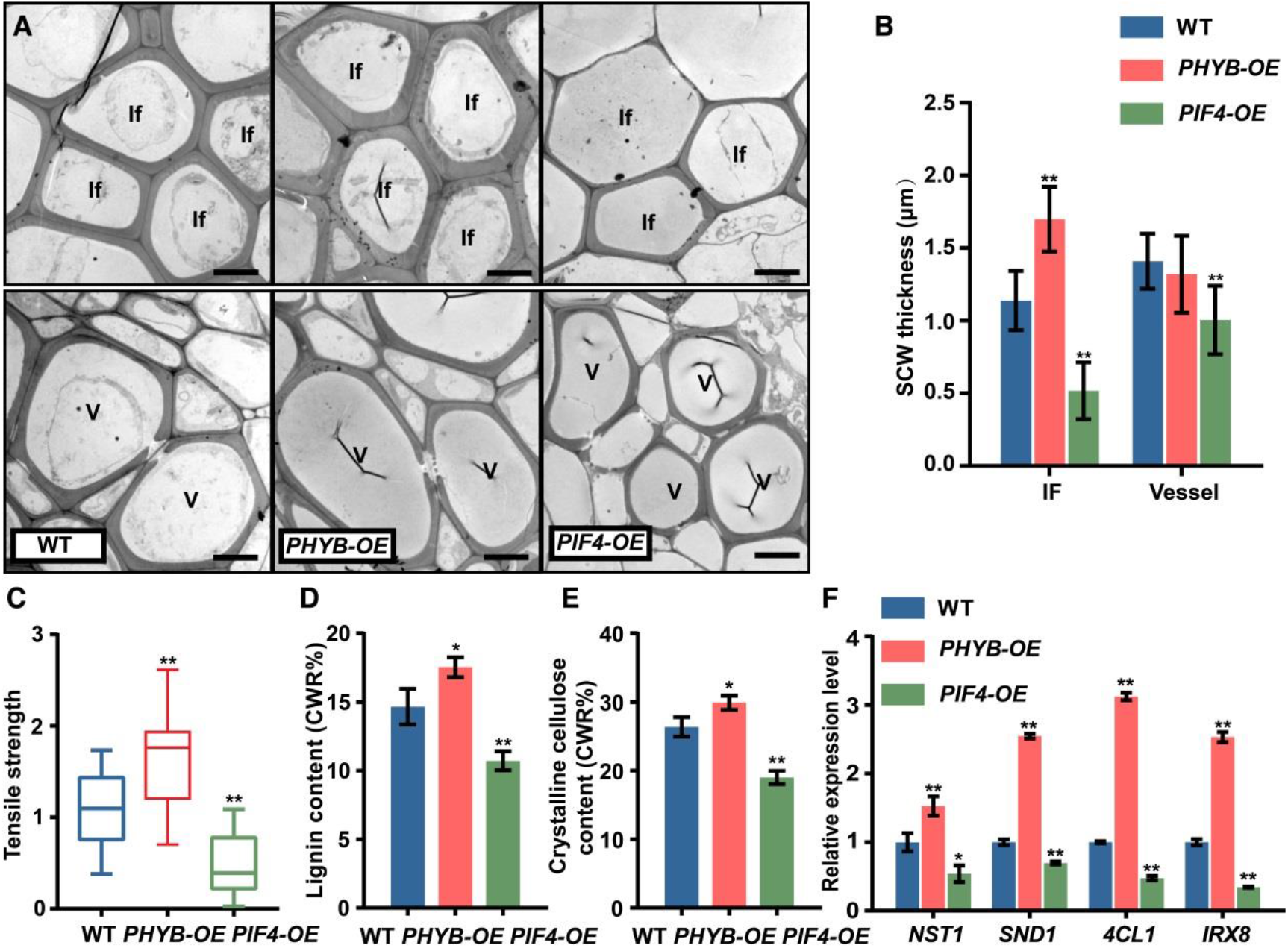
SCW phenotypes of *PHYB-OE* and *PIF4-OE* plants. **A,** Transmission electron micrographs of inflorescence stem cross-sections. If: Interfascicular fiber cell. V: Vessel cells. Scale bar = 5 μm. **B,** Measurements of SCW thickness of cells in (**A**). Data were from three biological replicates and more than 10 cells were measured per biological replicate. Student’s t test (**P < 0.01) was used for statistical analysis, mean ± SD. **C,** Tensile strength of inflorescence stems. Student’s t test (**P < 0.01) was used for statistical analysis, n = 15, mean ± SD. **D,** Lignin content in inflorescence stems. Student’s t test (*P < 0.05) was used for statistical analyses, n = 3, mean ± SD. **E,** Cellulose content in inflorescence stems. Student’s t test (**P < 0.01, *P < 0.05) was used for statistical analyses, n = 3, mean ± SD. **F,** Expression of SCW regulatory (*NST1* & *SND1*) and biosynthesis-related (*4CL1* & *IRX8*) genes in different genotypes grown under white light. Three biological repeats were performed. Student’s t test (**P < 0.01, *P < 0.05) was used for statistical analysis, mean ± SD.

### MYC2 links the *PHYB-PIF*s signal module to SCW thickening

As shown above, SCW thickening is genetically regulated by *PHYB* and *PIF*s and also conditionally modified by light R:FR ratio. Next, we examined whether the effect of PHYB and PIFs is dependent on the red-light signal. *phyB* and *pifq* mutants were grown under red light (high R:FR). The inflorescence stem of *phyB* mutant plants showed a significant decrease of the stem tensile strength, SCW thickening was severely impacted in fiber cells, but was essentially unchanged in vessel cells (**Figure 4, A-C**). Conversely, the *pifq* mutants displayed increased stem tensile strength, accumulated much thicker SCWs in xylary fiber cells and had a modest increase in vessel cells, compared to those in WT plants (**Figure 4, A-C**). Lignin content was decreased in the *phyB* mutant but increased in the *pifq* mutant (**Figure 4D**). Consistent with these findings, the expression of the SCW thickening genes (*NST1*, *SND1*, *4CL1* and *IRX8*) was downregulated in the *phyB* mutant but upregulated in the *pifq* mutant (**Figure 4E**). These data suggest that high R:FR facilitates SCW thickening through *PHYB*-enhanced SCW gene expression, while *PIF*s inhibit this process. Analysis of gene expression showed that *PHYB* and *PIF* quadruple members are both expressed in inflorescence stems, and among them *PIF4* and *PIF5* have the highest abundance (**Supplemental Figure S4A**). Also, *PIF4* promoter-GUS analysis indicated its expression in interfascicular fiber cells (**Supplemental Figure S4B**).

**Figure 4.**
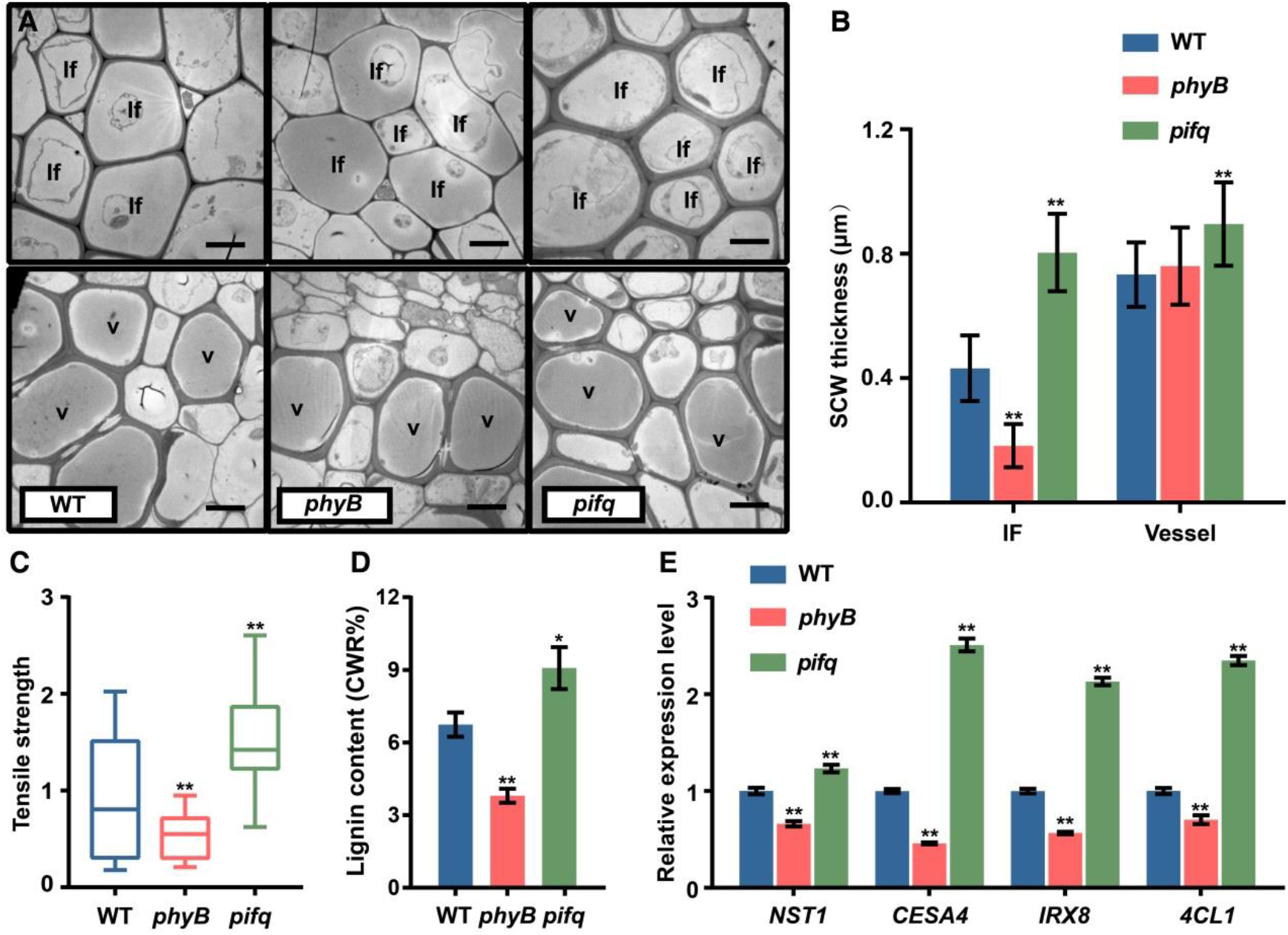
Red light signaling regulates SCW thickening dependent on *PHYB* and *PIF*s. **A,** The inflorescence stems of *phyB* and *pifq* mutants were grown under red light (high R:FR) and anatomically analyzed. Transmission electron micrographs of the inflorescence stem cross-sections. If: Interfascicular fiber cell. V: Vessel. Scale bar = 5 μm. **B,** Statistics of SCW thickness in (**A)**. Data were three biological replicates and more than 10 cells were measured per biological replicate. Student’s t test (**P < 0.01) was used for statistical analysis, mean ± SD. **C,** Tensile strength of the inflorescence stem. Student’s t test (**P < 0.01) was used for statistical analysis, n = 20, mean ± SD. **D,** Lignin content in the inflorescence stem. Student’s t test (**P < 0.01, *P < 0.05) was used for statistical analyses, n = 3, mean ± SD. **E,** Expression of SCW regulatory (*NST1*) and biosynthesis-related (*CESA4*, *4CL1* & *IRX8*) was measured by qRT-PCR analysis. Three biological replicates were performed. Student’s t test (**P < 0.01) was used for statistical analysis, mean ± SD.

Next, we dissected the molecular pathway that mediates the *PHYB-PIFs* signal to SCW thickening. Five-week-old WT plants at the stage of inflorescence stem elongation were moved from normal light to dark for 24 h in order to shut-down the light-induced gene expression, and then transferred to red light (high R:FR) conditions for 2 h. Stem samples were collected for RNA-sequencing (see **Supplemental Figure S5A**). Compared to 24 hours of dark status, a group of 2,203 differentially expressed genes (DEGs) were detected after red-light treatment, among which two thirds were up-regulated while one third were down-regulated (**Supplemental Table S1; Supplemental Figure S4B**). The regulated genes included red light responsive photopigment genes (*PSY*, *PORC*, *GUN5*) and target genes of PIFs (*PIL1*, *ATHB2*, *BBX28*)(Zhang et al., 2013; Toledo-Ortiz et al., 2014), indicative of the red-light signaling effectiveness. It was noticed that the expression of cell expansion genes (*XTH22*, *XTH27*, *XTH30*, *EXPA1*) was inhibited, but the SCW thickening-related regulatory genes, including *NST1* and *MYC2*, were up-regulated in response to red light (**Supplemental Figure S5C**)(Matsui et al., 2005; Claisse et al., 2007; Goh et al., 2012; Taylor-Teeples et al., 2015). Interestingly, expression of *VND6*/*VND7* which control vessel SCW thickening was not induced by red light (**Supplemental Table S1**). The red-light induction of *NST1*, *MYC2* and other key genes for SCW thickening was further confirmed in RT-qPCR analysis (**Supplemental Figure S6**). Conversely, when exposed to far-red light conditions, *MYC2* and *NST1* expression was inhibited in WT plants, however, the inhibition was reduced in *phyB* and *pifq* mutants (**Supplemental Figure S7**). These results suggest that the expression of *MYC2* is induced by red light and inhibited by far-red light and regulated through PHYB and PIFs.

### PIF4 affects MYC2 stability and inhibits its transcriptional activity

It is known that MYC2 binds to the *NST1* promoter to regulate SCW thickening(Zhang et al., 2018a). To analyze how PIFs affect MYC2 function in SCW thickening, we conducted a dual-luciferase (LUC) reporter assay in *Arabidopsis* protoplasts. We co-expressed PIFs with MYC2 to check the MYC2 transcriptional activity on the *NST1* promoter. MYC2, but neither PIF4 nor PIF5, were able to activate the *NST1* promoter (**Figure 5, A and B**). When MYC2 was co-expressed with either PIF4 or PIF5, the MYC2 activity on the *NST1* promoter was reduced (**Figure 5B**), suggesting that the MYC2 transcriptional activity is repressed by either PIF4 or PIF5. Studies have shown an interaction between PIFs and MYC2 to contribute to apical hook formation and plant defense(Zhang et al., 2018b; Zhao et al., 2021). Our yeast two-hybrid assays verified the PIF4 interaction with MYC2 is likely through the PIF4 N-terminal region (**Supplemental Figure S8**).

**Figure 5.**
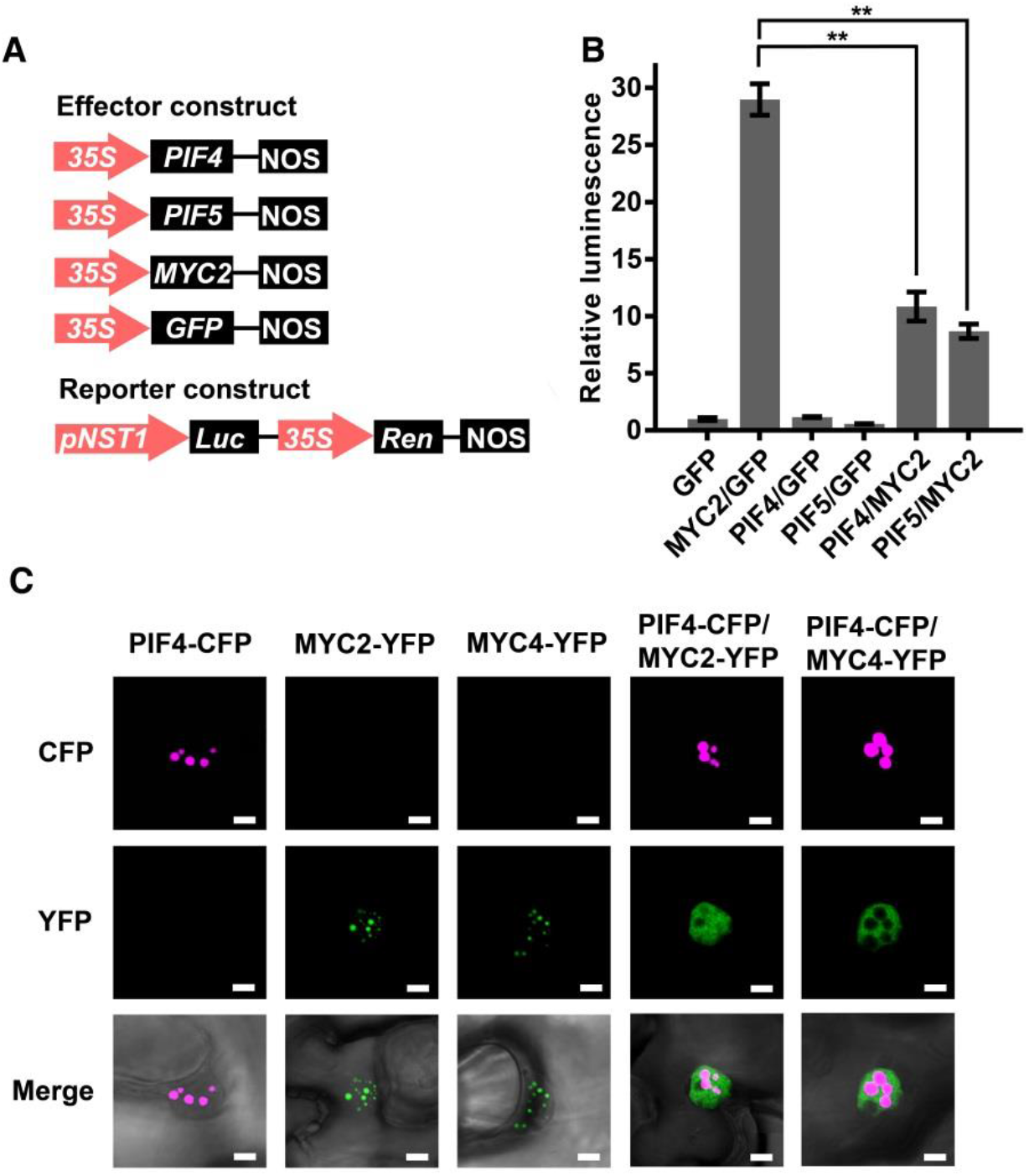
PIF4 represses the MYC2 transcriptional activity. **A,** Schematic representation of the *NST1* promoter-driven dual-LUC reporter gene and three effector gene constructs. 35S promoter, *NST1* promoter (−1 to −3711 bp from ATG), renilla luciferase (REN), firefly luciferase (LUC) are indicated in reporter constructs. In effector constructs, PIF4, PIF5 and MYC2 are driven by the 35S promoter. **B,** PIF4/PIF5 inhibit MYC2 activation of *NST1* promoter. *Arabidopsis* protoplasts were transfected with the reporter constructs in combination with different effector consrtructs. After transfection, the protoplasts were kept in dark for 16 h. Relative luminescence was normalized to the protoplast transformed with reporter and empty effector (GFP). Student’s t test (**P < 0.01) was used for statistical analysis, n = 3, mean ± SD. **C,** Subcellular localization of PIF4 and MYC2/MYC4. Constructs of PIF4-CFP, MYC2-YFP and MYC4-YFP were transferred to tobacco leaves, separately or together, by Agro-infiltration. Then the tobacco leaves were kept in dark for 12 h before fluorescence observation. Scale bar = 5 μm

Then, the subcellular localization of MYC2, its homolog MYC4 and PIF4 was examined. Tobacco leaf cells were employed to express MYC2/MYC4-YFP and PIF4-CFP proteins and then treated with dark condition. When expressed separately, PIF4 and MYC2/MYC4 proteins were localized in the nucleus as distinct dots. However, when PIF4 and MYC2/MYC4 were co-expressed together, the nuclear localization of PIF4 was unchanged, whereas MYC2 and MYC4 became distributed throughout the nucleus (**Figure 5B**). These results suggest that PIF4 affects MYC2/MYC4 nuclear localization and/or stability.

Next, MYC2 protein stability was tested in *planta*. MYC2-YFP was expressed in wild-type and *pifq* mutant plants. Immunoblot analysis showed that the MYC2 abundance in inflorescence stems was decreased in both dark and in far-red light conditions, while MYC2 is more stable in the *pifq* mutant plants compared to the WT background (**Figure 6, A and B**). Furthermore, the MYC2-YFP fluorescence signal in root tip cells showed a substantial decrease in the WT background after dark treatment, but the reduction was attenuated in the *pifq* mutant background (**Figure 6C**). These results suggest that PIFs affect MYC2 protein stability.

**Figure 6.**
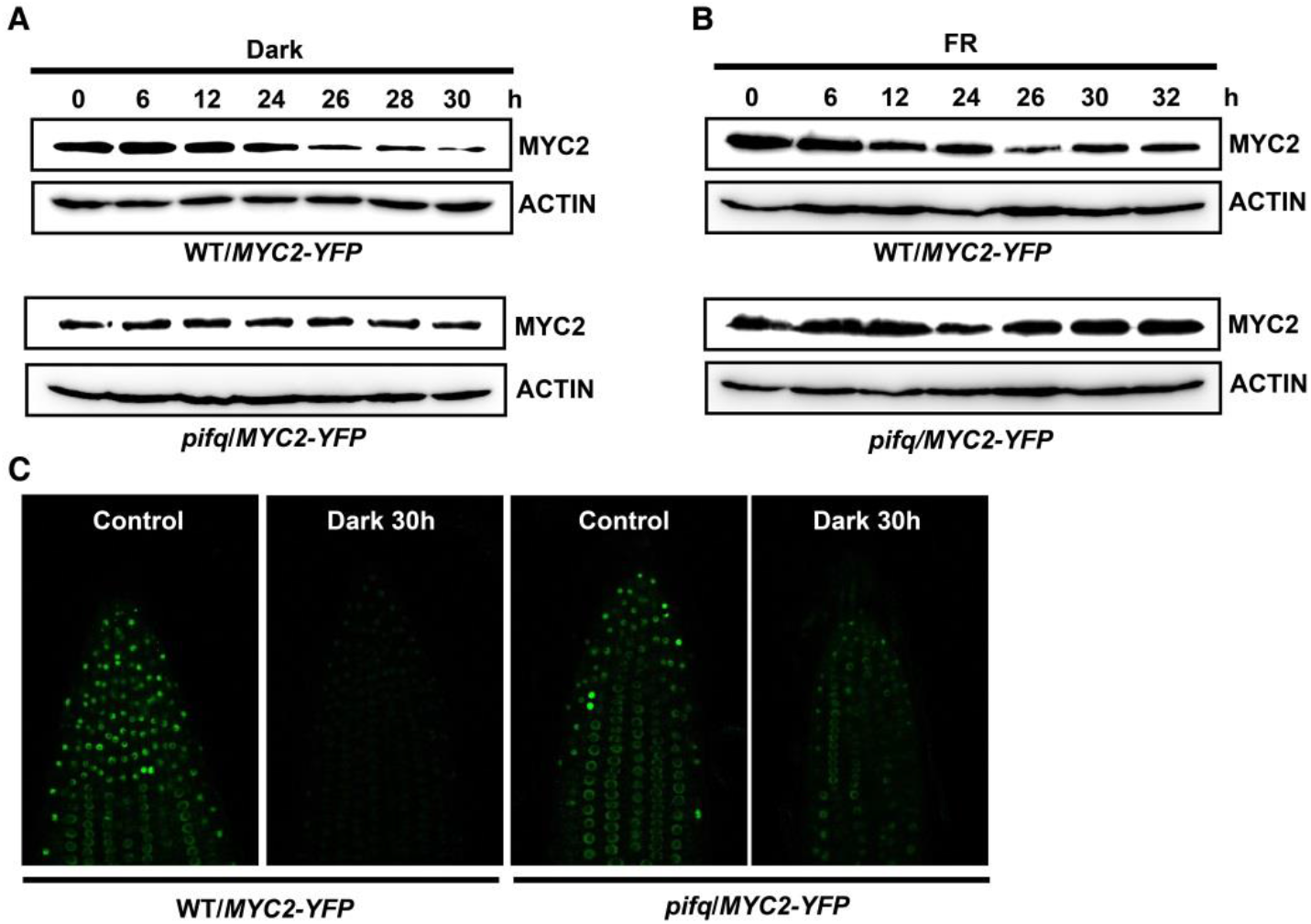
PIF4 affects MYC2 stability in dark and FR light. **A,** Immunoblot of MYC2-YFP and ACTIN in transgenic plants. Seedlings were grown in WL condition for 10 days and then treated with dark condition. Similar results were observed in three independent experiments. **B,** Immunoblot of MYC2-YFP and ACTIN in transgenic plants. Seedlings were grown in WL condition for 10 days and then treated with far-red light. Similar results were observed in three independent experiments. **C,** Fluorescence signals in seedling roots. Seedlings were grown in WL condition for 4 days and then treated with/without dark for 30 h.

### *MYC2*/*MYC4* act downstream of *PHYB* and *PIF*s in a genetic pathway to regulate stem SCW thickening

Next, *myc2myc4* double mutants were crossed with *pifq* and *phyB* mutants to test whether *MYC2* lies in the same genetic pathway to *PHYB* and *PIF*s in the regulation of SCW thickening. The *pifq* phenotype of a shorter inflorescence stem relative to WT (**Figure 2, A and B**) was restored in the sextuple *pifqmyc2myc4* mutant under white light (**Figure 7, A and B**). Consistently, both fiber cell thickness (**Figure 7, C and D**) and expression of the SCW thickening related genes such as *NST1*, *4CL1*, and *IRX8* were decreased in *pifqmyc2myc4* plants compared with *pifq* (**Figure 7, E and F**).

**Figure 7.**
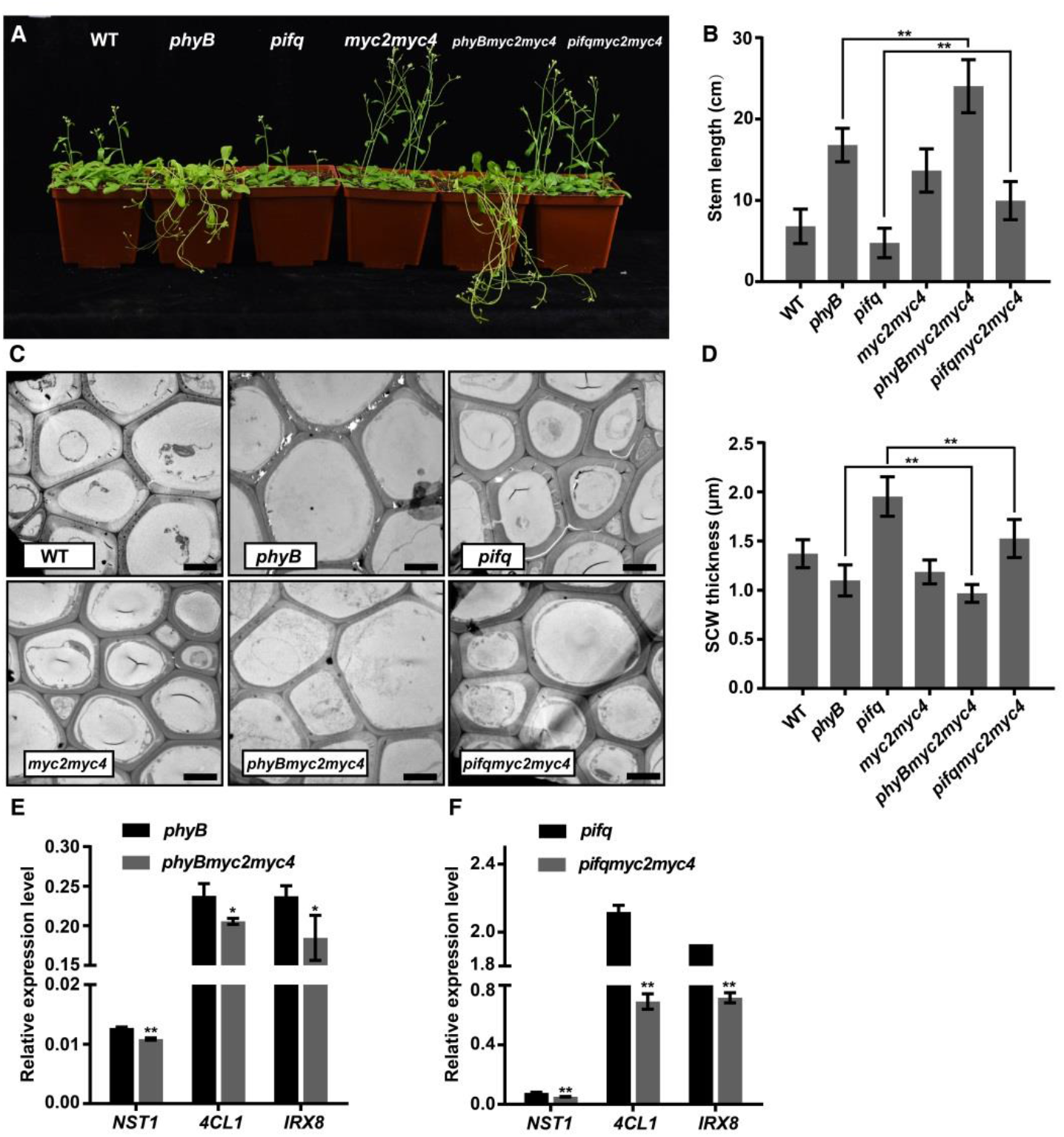
*MYC2*/*MYC4* genetically interact with *PIFs*. **A,** Mutant plants (*phyB*, *pifq*, *myc2myc4*, *phyBmyc2myc4*, *pifqmyc2myc4*) were grown in white light until 4-weeks old. Scale bar = 5 cm. **B,** Inflorescence stem length in various mutants. Student’s t test (**P < 0.01) was used for statistical analysis, n > 10, mean ± SD. **C,** Transmission electron micrographs of stem cross sections showing interfascicular fiber cells. Scale bar = 5 μm. **D,** Statistics of SCW thickness in interfascicular fiber cells in (**C**). Data were collected from three biological replicates and more than 10 cells were measured per biological replicate. Student’s t test (**P < 0.01) was used for statistical analysis, mean ± SD. **E, F,** Expression of the key SCW regulatory (*NST1*) and biosynthesis-related (*4CL1* & *IRX8*) genes in mutant plants. Three biological replicates were performed. Student’s t test (**P < 0.01, *P < 0.05) was used for statistical analysis, mean ± SD.

Furthermore, rosette leaf growth and hypocotyl elongation, which were inhibited in *pifq*(Leivar and Quail, 2011), was partly rescued in *pifqmyc2myc4* plants (**Supplemental Figure S9, A and D and E**). These results indicate that *MYC2*/*MYC4* are genetically downstream of the *PHYB*-*PIFs* signaling module. Nevertheless, the faster elongating-stem and thinner-SCW phenotypes of *phyB* mutants were enhanced in *phyBmyc2myc4* triple mutants (**Figure 7, A-D**), and the SCW regulatory and biosynthesis-related gene expression was slightly down-regulated in triple mutants compared with *phyB* (**Figure 7E**). Additionally, *phyBmyc2myc4* mutants exhibited greater hypocotyl elongation than *phyB* seedlings (**Supplemental Figure S9, B and C**). Together, these data indicate that the *PHYB*-*PIFs* signaling module regulates SCW thickening in the inflorescence stem of *Arabidopsis*. *MYC2* acts downstream of *PIF*s to modify the transcription networks of SCW thickening to tune plant stem growth.

## Discussion

### Shaded light promotes cell elongation as well as inhibits SCW thickening in inflorescence stem of *Arabidopsis*

In the xylem of the plant vascular system, fiber and vessel cells develop a thickened SCW to provide mechanical support and facilitate water transport(Lucas et al., 2013). SCW thickening in xylem cells is regulated by growth and developmental cues as well as environmental conditions through a combination of hormone and mechano-sensing signals(Didi et al., 2015; McCahill and Hazen, 2019; Hori et al., 2020). Plants grow longer petioles and inflorescences with weak stem strength and thinner SCW under shade conditions(Kozuka et al., 2010; Casal, 2012; Wu et al., 2017), but the mechanisms of how the shaded light modulates SCW thickening and mechanical properties are largely unknown.

A characteristic feature of a shaded light condition is the decreased red:far-red (R:FR) light ratio(Hersch et al., 2014). Using the *Arabidopsis* inflorescence stem as a model, we examined the effect of the R:FR ratio on the SCW thickening of interfascicular fibers and vessel cells, the major cell types with SCW. A low ratio of R:FR light induced rapid elongation of the inflorescence stem resulting in a lodging phenotype (**Figure 1, A and B**), consistent with the reported shaded light effect on stem growth(Casal, 2012), indicating that the low ratio of red:far-red light can be used as an experimental proxy to simulate shaded light conditions. Anatomical analyses reveal that the shaded light results in plants with a thinner SCW in stem fiber cells, and weaker mechanical strength of the inflorescence stem (**Figure 1, A and C-E**). Expression of SCW regulatory and biosynthesis-related genes is down-regulated (**Supplemental Figure S1**). Many plants respond to low R:FR ratio light with the shade avoidance syndrome (SAS), displaying a series of well described morphological changes such as enhanced elongation of hypocotyl, internode and petiole(Liu et al., 2021). In contrast, the cellular and molecular mechanisms underpinning these morphological phenotypes to a low R:FR ratio light have not been well defined. Plant morphogenesis and body structure is defined by the assembly and architecture of the cell wall(Huang et al., 2018). Our study shows that SCW thickening, and cell elongation are coordinated and tightly regulated by shaded light conditions.

### *PHYB*-*PIFs* signaling module mediates regulation of SCW thickening in inflorescence stem of Arabidopsis

The *PHYB*-*PIFs* signaling module plays a primary role in plant SAS and morphogenesis through PHYB-PIFs-controlled gene expression(Reed et al., 1993; Pham et al., 2018). Shaded light inactivates PHYB and subsequently diminishes its action on the degradation of PIFs, leading to regulation of the downstream genes to modulate plant growth(Lorrain et al., 2008; Jia et al., 2020). Whether PHYB and PIFs are directly involved in the shaded light inhibition of SCW thickening is unresolved. To answer this question, PHYB and PIFs function on cell wall formation of inflorescence stem was examined. *phyB* and *pifq* mutants have contrasting phenotypes of SCW thickening and mechanical strength properties in inflorescence stem (**Figure 2 and 3**), indicating that the SCW thickening process is positively regulated by PHYB but negatively regulated by PIFs.

In *Arabidopsis* inflorescence stems, SCW formation in the fiber and vessel cells are initiated under the control of the master TF switches of *NST1*/*SND1*(Zhong et al., 2006; Mitsuda et al., 2007) and *VND6*/*VND7*(Yamaguchi et al., 2010), respectively. *NST1*/*SND1* and *VND6*/*VND7* regulate several downstream genes in controlling SCW thickening(Taylor-Teeples et al., 2015) through highly cell-type specific spatio-temporal regulation modulated through distinct regulatory signals(Zhong et al., 2006; Zhong et al., 2008). In this study, low R:FR ratio results in thinner SCW in interfascicular fiber cells of inflorescence stems while the vessel SCW were largely unaffected (**Figure 1, D and E**). Genetic evidence reveals that the PHYB-PIFs interaction mediates the signaling underlying the low R:FR ratio inhibition of SCW thickening in fiber cells but not in vessel cells (**Figure 2**). Interestingly, the blue light signal dramatically affects SCW thickening in fiber cells with little effect on vessel cells(Zhang et al., 2018a). Information from RNA sequencing indicated that expression of *VND6*/*VND7* does not respond to red-light induction (**Supplemental Table S1**). Therefore, light-induced SCW thickening appears to be through the transcriptional network directed by *NST1* in fiber rather than xylem cells.

### Shaded light inhibits SCW thickening through PIF4 inactivation of MYC2 activity

MYC2 interconnects a variety of environmental and developmental responses(Yadav et al., 2005; Chen et al., 2012; Song et al., 2014). It is regulated at both transcriptional and post-translational levels under various circumstances. Blue light induces the expression of *MYC2* and *MYC4* and subsequently activates *NST1* to promote SCW thickening(Zhang et al., 2018a). MYC2 protein is targeted for degradation during hormone responses(Jung et al., 2015; Chico et al., 2020). In addition, MYC2 protein is known to be destabilized under either dark or shade conditions, while red light stabilizes MYC2 in a PHYB-dependent manner(Chico et al., 2014), indicating MYC2 plays a role in the light signaling pathway.

We demonstrate that *MYC2,* as well as the downstream expression of the SCW thickening genes, are enhanced upon high R:FR light induction and reduced after a far-red light treatment (**Supplemental Figure S5-7**). Meanwhile, MYC2 protein is destabilized under far-red light and dark conditions in WT (**Figure 6, A and B**) but more stable in the *pifq* mutant under the same treatments (**Figure 6, A and B**). Therefore, we conclude that the *PHYB-PIFs* signaling module affects MYC2 protein stability.

PIFs interact with MYC2 and such interaction plays a role in regulating the *NST1* promoter transcriptional activity (**Figure 5, A and B**). MYC2/MYC4 activates *NST1* transcription by binding its promoter, but PIF4 is inactive to *NST1* transcription. MYC2/MYC4 and PIF4 are localized in nuclei as distinct dots (**Figure 5C**), similar to that previously reported(Withers et al., 2012; Luo et al., 2014). However, when PIF4 and MYC2/MYC4 were co-expressed in the dark, PIF4 localization was unchanged, whereas the dotted localization of MYC2 and MYC4 became distributed throughout the nucleus (**Figure 5C**). This suggests that interaction of PIF4 with MYC2/MYC4 displaced MYC2 localization in the nucleus. It is known that low R:FR light condition prevents PHYB from entering the nucleus and PIF4 becomes stable to interact with other proteins(Chico et al., 2014; Leivar and Monte, 2014). In our study, PIF4 interacts with MYC2/MYC4 under shaded light conditions to either prevent MYC2/MYC4 binding to the *NST1* promoter or induces MYC2 protein degradation, leading to the inhibition of MYC2/MYC4 transcriptional activity. Given that PIF proteins have been shown to be regulators of COP1 ubiquitination activity(Jang et al., 2010) and that MYC2 protein abundance in *cop1* mutants is at a sustained high level(Chico et al., 2014), it is possible that PIFs enhance MYC2/MYC4 polyubiquitination by COP1 to promote its degradation.

The interaction between PIF4 and MYC2/MYC4 in regulating SCW thickening was further confirmed genetically by crossing *myc2myc4* with *pifq* and *phyB* mutants. By analyzing the SCW phenotypes in inflorescence stem elongation growth (**Figure 7**), we concluded that *MYC2* and MYC4 function genetically downstream of *PHYB* and *PIFs* to regulate SCW thickening in *Arabidopsis*. A recent study showed that overexpression of *MYC2* in a *phyB* background partially suppressed its long hypocotyl phenotype(Ortigosa et al., 2020), suggesting a genetic interaction of *PHYB* and *MYC2* in hypocotyl elongation.

To summarize the findings of this study, we propose a model for the shaded light inhibition of SCW thickening (**Figure 8**): Under normal light conditions (high R:FR ratio), PHYB undergoes a light-dependent conformational change, which relieves the interaction between chromophore-attachment domains and PAS-related domain (PRD), and the nuclear-localization signal in the PRD is unmasked, enabling PHYB to translocate to the nucleus and promote phosphorylation and degradation of PIF proteins(Chen et al., 2005), thus resulting in MYC2/MYC4 stabilization and *NST1* transcriptional activation to promote SCW thickening. Upon perception of shaded light (low R:FR ratio), PHYB undergoes dark reversion from Pfr to its inactivated Pr form, and returns to its cytoplasmic location since its nuclear-localization signal is masked, allowing accumulation of PIFs in the nucleus where they interact with MYC2/MYC4. Thus, the MYC2 activity on *NST1* transcription is inhibited, leading to suppression of the SCW thickening (**Figure 8**). The study reveals a mechanism by which shaded light inhibits SCW thickening.

**Figure 8.**
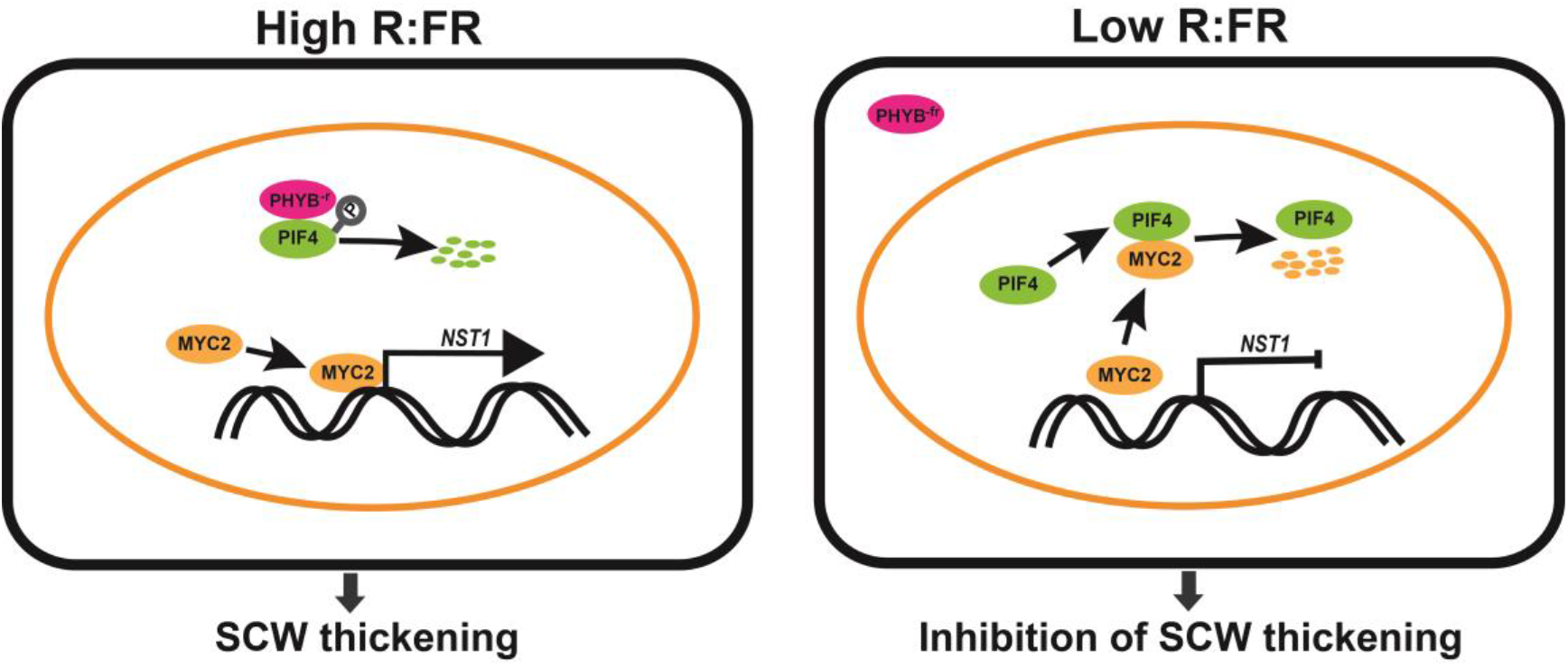
A proposed model of shaded light regulation of the SCW thickening. White light enhances SCW thickening in fiber cells of inflorescence stem. Under white light (high R:FR condition), PHYB is activated to its Pfr form which enters the nucleus to inhibit PIF activity, MYC2 is available to bind to the *NST1* promoter to activate the *NST1*-direct SCW thickening process. Under shaded light (low R:FR condition), PHYB reverts to its inactive Pr form. PHYB cannot enter the nucleus and PIF proteins interact with MYC2 and displaces its binding to the *NST1* promoter. Thus, the *NST1*-directed SCW thickening process is suppressed.

Since regulation of SCW thickening is coordinated with cell elongation, our finding that light signaling modules regulate SCW thickening has important implications for both our understanding of plant growth and development and proffers a pathway to begin to modify the mechanical properties of secondary cell walls that are crucial to both forest (wood and wood product properties) and agricultural (crop improvement through, for example, prevention of lodging) industries.

## Materials and methods

### Plant materials and growth conditions

The *Arabidopsis* wild-type (WT) ecotype used in this study is in a Columbia-0 (Col-0) background. The *phyB, pifq, PIF4-OE, PHYB-OE* and *myc2myc4* plants were generated as described previously (Wang et al., 2010; Jia et al., 2014; Ma et al., 2016; Zhang et al., 2018a). The *phyBmyc2myc4* and *pifqmyc2myc4* mutant plants were generated by crossing *myc2myc4* with either *phyB* or *pifq* and genotyped using primers listed in **Appendix Table S2**. To generate WT/*35S:MYC2-YFP* and *pifq*/*35S:MYC2-YFP* plants, *MYC2* coding sequence was cloned into a pHB-X-YFP vector and transferred to either WT or *pifq Arabidopsis* following the floral dip method with the *Agrobacterium tumefaciens* strain GV3101(Clough and Bent, 1998). Plants were grown in a phytotron with a light (fluorescent lamp, 80μE/s*m^2^) /dark cycle of 16 h/8 h at 22°C. For light treatment of inflorescence stem growth, plants were grown under white light until bolting and transferred to various light conditions. A red-light condition was achieved with a light-emitting diode (LED) light incubator (Percival E30LED; 30 μE/s*m^2^). White light (WL; high R:FR) was provided by a WL LED with a R:FR ratio of 13 (Qiding technology, 67 μE/s*m^2^). WL+FR (low R:FR) treatment was achieved using supplementary far-red (FR) LEDs (Qiding technology, 47 μE/s*m^2^) to a R:FR ratio of 0.066. FR light was provided by FR LEDs (Qiding technology, 47 μE/s*m^2^). All light parameters were measured with an ILT1400 Radiometer Photometer and an Ocean Optics HR2000+CG spectrophotometer.

### Paraffin sections

Paraffin sections were performed as previously described (Zhang et al., 2018a). Briefly, the basal part of the inflorescence stem was cut into a 5-mm segment and fixed in FAA (Formalin-Aceto-Alcohol) solution under vacuum, stored at 4°C overnight, dehydrated in a graded ethanol series, and embedded into paraffin. Samples were sectioned to 10 μm thickness using a Leica RM2235 rotary microtome. The sections were stained with Toluidine blue and observed under a light microscope (Olympus, BX53).

### Cell wall thickness analysis

The basal part of the inflorescence stem was cut into 2 mm segments, fixed in 3% (v/v) paraformaldehyde and 0.5% (v/v) glutaraldehyde in PBS (0.1 M, pH 7.4), dehydrated in a graded ethanol series, embedded in Epon812 and sectioned. The samples were stained with 2% (w/v) uranyl acetate and lead citrate and observed under a transmission electron microscope (Hitachi H-7650) as previously described (Zhao et al., 2014). SCW thickness was measured with ImageJ software.

### Fiber cell length measurement

The basal part of the inflorescence stem was cut into a 2 cm length and disaggregated by submerging in glacial acetic acid / 30% hydrogen peroxide (v/v 1:1) solution at 60 ℃ overnight, stained with 1% safranine (w/v) for 10 min, and photographed under a light microscope (Olympus, BX53). The fiber cell length was measured with ImageJ software.

### RNA extraction and quantitative RT-PCR analysis

Total RNA was extracted from different tissues of *Arabidopsis* plants using the E.Z.N.A. Plant RNA Kit (Omega, R6827-02). The first-strand cDNA was synthesized using TransScript One-Step gDNA Removal and cDNA Synthesis SuperMix (TransGen Biotech, AT311-03) for quantitative real-time PCR (qRT-PCR) analysis of transcript abundance. qRT-PCR was performed using SYBR Green (TransStart Tip Green qPCR supermix) with a QuantStudio 3 Real-Time PCR System (Applied Biosystems). Gene expression was normalized using *ACT2* as an internal control.

### Analysis of stem tensile strength and cell wall components

The basal part of the inflorescence stem was used for tensile strength measurement as previously described (Zhang et al., 2018a). Relative tensile strength was normalized against WT. For analysis of cell wall components, the basal part of inflorescence stem was collected and analysed as described by (Xi et al., 2017). Collected stems were ground to a fine powder in liquid nitrogen, Alcohol-insoluble residue (AIR) was obtained by successively washing the powder with 70% (v/v) ethanol, chloroform/methanol (1:1 v/v), and acetone (Pettolino et al., 2012). After de-starching, AIR was washed with water and acetone, and dried for lignin and crystalline cellulose content determination.

### Immunoblotting

Proteins extracted from seedling samples were separated with 10% (w/v) SDS-PAGE gels and blotted onto polyvinylidene fluoride membrane (Bio-rad). Protein blots were then analyzed using either anti-GFP/Myc (1:2000 dilution; Abmart) or anti-ACTIN (1:2000 dilution; Abmart) monoclonal antibodies followed by horseradish peroxidase (HRP)-conjugated goat-anti-mouse antibodies (1:5000 dilution, Thermo Fisher). Blots were developed in a Tanon Imaging System (Tanon 5200CE) using ECL Western Blotting Substrate (Tanon, 180-501).

### Protein subcellular localization

For protein subcellular localization analyses, the MYC2/MYC4 and PIF4 coding sequences were PCR-amplified with the proof-reading enzyme PHANTA (Vazyme, P520) from cDNA and cloned into a pHB-X-YFP and a pHB-X-CFP vector in frame with the YFP/CFP (Luo et al., 2014), respectively, and then the constructs were Agro-infiltrated into *Nicotiana benthamiana* leaves according to the method of (Gui et al., 2016). After incubation for 48 h in dark, abaxial epidermal cells of the leaf were observed under a confocal microscope (Leica TSC SP8 STED 3X).

### Yeast two-hybrid assay

The coding sequence of MYC2 PCR-amplified from cDNA using PHANTA was fused with GAL4 DNA-binding domain (BD) of the bait vector pGBKT7 (Clontech). N-terminal containing TAD and C-terminal containing bHLH of PIF4 were PCR-amplified and fused with GAL4 activation domain (AD) of the pray vector pGADT7 (Clontech). Bait and pray vectors were co-transformed in to the Y2H Gold yeast strain, and then grown on SD-Trp-Leu and SD-Trp-Leu-His plates (Clontech).

### PIFs and MYC2 transcriptional activity assay

The interaction of MYC2 and PIF4 activity was assayed using a dual-LUC reporter assay system (Promega) through *Arabidopsis* protoplast transfection. The PIF4/ PIF5/ MYC2 and GFP coding sequences were cloned into the pA7 vector under the control of the 35S promoter and used as an effector. The *NST1* (−1 to −3711 bp from ATG) promoter sequence was cloned into *pGreenII 0800-LUC* vector upstream and in frame with the luciferase (LUC) gene and was used as a reporter. Renilla luciferase (REN) gene in a pGreenII0800-LUC vector was used as an internal control. Protoplasts from *Arabidopsis* mesophyll cells were isolated and transformed as previously described (Zhang et al., 2018a).

### Promoter GUS activity analysis

The *PIF4* promoter (−1 to −4312 bp from ATG) was cloned into a pORE-R2 vector (Fang et al., 2021) to drive GUS expression, then the promoter constructs were transformed into wild-type *Arabidopsis* following the floral dip method (Clough and Bent, 1998). Transgenic plants were selected on MS medium containing 50 μg/ml hygromycin and then genotyped. Positive T2 transgenic plants were used to analyze the GUS staining activity. GUS staining was conducted according to (Gui et al., 2016).

### Accession Numbers

Sequence data from this article can be found at https://www.arabidopsis.org with the following accession numbers: *PHYB* (AT2G18790), *PIF1* (AT2G20180), *PIF3* (AT1G09530), *PIF4* (AT2G43010), *PIF5* (AT3G59060), *MYC2* (AT1G32640), *MYC4* (AT4G17880), *NST1* (AT2G46770), *SND1* (AT1G32770), *VND6* (AT5G62380), *VND7* (AT1G71930), *MYB103* (AT1G63910), *CESA4* (AT5G44030), *IRX8* (AT5G54690), *4CL1* (AT1G51680), *LAC4* (AT2G38080), *PER64* (AT5G42180), *F5H* (AT4G36220), *PIL1* (AT2G46970), *ATHB2* (AT4G16780), *BBX28* (AT4G27310), *PSY* (AT5G17230), *PORC* (AT1G03630), *GUN5* (AT5G13630), *XTH27* (AT2G01850), *XTH22* (AT5G57560), *XTH30* (AT1G32170), *EXPA1* (AT1G69530) and *ACT2* (AT3G18780).

## Supplemental data

The following materials are available in the online version of this article.

**Supplemental Figure S1.** Expression of SCW-related genes under different light conditions.

**Supplemental Figure S2.** PHYB and PIFs affect inflorescence stem properties.

**Supplemental Figure S3.** Phenotypes of PHYB-OE and PIF4-OE plants.

**Supplemental Figure S4.** Expression pattern of phyB and PIF genes.

**Supplemental Figure S5.** Transcriptional analysis of Arabidopsis inflorescence stem treated with red light.

**Supplemental Figure S6.** Expression of MYC2 and SCW formation-related genes is induced by red light.

**Supplemental Figure S7.** Far-red light inhibition MYC2 expression is dependent on PHYB and PIFs.

**Supplemental Figure S8.** PIF4 physically interacts with MYC2.

**Supplemental Figure S9.** myc2myc4 mutation rescued the phenotype of pifq mutant.

**Supplemental Table S1.** RNA-seq data of red light induced genes in 5 week old inflorescence stems.

**Supplemental Tabel S2.** Primer sequences used in this study.

## Acknowledgment

We thank Dr. Daoxin Xie (Tsinghua University) for providing *myc2 myc4* mutants. We thank Ni Fan and Dr. Xiaoshu Gao for help on confocal microscoping. We thank Xiaoyan Gao, Zhiping Zhang, Jiqin Li and Ling Ge for assistance with transmission electron microscopy. This work was supported by the National Natural Science Foundation of China (Grant No. 31630014) and the Chinese Academy of Sciences (Grant No. XDB27020104). MSD and AB would like to acknowledge the support of funds from La Trobe University and the Chinese national and provincial governments to the Sino-Australia Cell Wall Research Centre, Zhejiang Agriculture and Forestry University (ZAFU).

## Author contributions

F.L., Q.Z., H.L., H.Y. and L.L. designed the research; F.L., Q.Z., performed the experiments; F.L., Q.Z., H.L., H.Y., M.S.D., A.B. and L.L. analyzed the data; F.L., Q.Z, M.S.D. A.B. and L.L. wrote the paper. All authors read and approved the article.

